# Decoding the Microbiome-Disease Axis with Interpretable Graph Neural Networks

**DOI:** 10.1101/2025.06.23.659608

**Authors:** Vladimir A. Ivanov, Wyatt H. Hartman, Mohammad Soheilypour

## Abstract

The human gut microbiome is a complex ecosystem whose disruption is implicated in a wide spectrum of diseases, yet translating microbiome research into actionable therapeutics is hindered by a critical trade-off: existing models either prioritize predictive accuracy at the expense of interpretability or sacrifice performance for mechanistic insight, limiting their ability to pinpoint specific disease-driving microbial interactions and taxa. To address this, we introduce Graph neural network for Interpretable Microbiome (GIM), a graph neural network framework that integrates minimally processed taxonomic metadata as sparse node embeddings within an unweighted complete graph, enabling direct modeling of high-order microbial interactions through message passing. GIM achieves state-of-the-art classification performance on microbiome-disease prediction tasks (e.g., healthy vs. allergic states) while generating fine-grained, experimentally validated attributions at the level of taxonomic ranks, driver microbes, and putative microbe-to-microbe interactions. By bridging the gap between predictive accuracy and biological interpretability, GIM overcomes a key limitation in current approaches, offering a unified framework to both predict dysbiosis-associated disease states and identify actionable microbial targets for therapeutic intervention. This dual capability represents a critical advance toward precision microbiome engineering and scalable hypothesis generation in translational microbiome research.

## 1 Introduction

The human gut microbiome has emerged as a critical regulator of host physiology and is increasingly implicated in a broad spectrum of diseases De Vos et al. [2022]. Dysbiosis of gut microbial communities has been linked to metabolic disorders such as obesity and type 2 diabetes Fan and Pedersen [2021], autoimmune conditions such as rheumatoid arthritis Zhang et al. [2020] neurological disorders such as Parkinson’s disease and autism spectrum disorder Sorboni et al. [2022], and chronic inflammatory diseases such as inflammatory bowel disease (IBD) Glassner et al. [2020]. These widespread associations suggest that the gut microbiome may function not only as a biomarker of disease-associated states but also as a modifiable factor with therapeutic potential. This has driven significant interest in therapeutic strategies aimed at reshaping the microbiome, including probiotics, prebiotics, dietary interventions, and live biotherapeutic products, which seek to restore microbial diversity and promote beneficial metabolic functions Hitch et al. [2022]. However, translating these findings into effective therapeutics requires a mechanistic understanding of the complex microbial interactions that drive disease processes.

Despite clear associations between microbiome composition and disease outcomes, modeling microbiome data remains challenging. The human gut microbiome is a high-dimensional, compositional dataset: a typical sample may contain hundreds of bacterial taxa whose abundances vary across individuals Lin and Peddada [2020]. Traditional analytical approaches in microbiome research often focus on identifying individual differentially abundant taxa or diversity measures between health and disease groups Lutz et al. [2022]. While these statistical comparisons (and classical machine learning models such as random forests or logistic regression) have yielded important candidate “microbial biomarkers,” they largely ignore the complex network of interactions within the microbial community. In reality, microbes do not act in isolation – they engage in cooperative and competitive interactions (for nutrients, niches, signaling) that can collectively govern community stability and functional output Wu et al. [2020], Collins et al. [2023]. Notably, disruptions in certain keystone species may have cascading effects on the broader community Cho and Blaser [2012], and groups of microbes may exhibit emergent properties (such as metabolic co-dependence) relevant to disease processes Ackermann et al. [2023]. Failing to account for these inter-microbial relationships can limit our understanding of dysbiosis and reduce the accuracy of predictive models. There is thus a compelling need for computational frameworks that move beyond treating the microbiome as a mere list of independent features, toward modeling it as an interconnected ecosystem.

A fundamental challenge in this domain is the precise reconstruction of these interaction networks (“wiring diagrams”) and the identification of driver microbes capable of modulating community states toward health or disease trajectories Liu [2023], Tan et al. [2024], Wang et al. [2021]. Limitations of traditional microbiome modeling approaches have led to the adoption of deep learning (DL) methods to better capture the complexity inherent in microbiome data Baranwal et al. [2022], Ghannam and Techtmann [2021]. Grounded in the universal approximation theorem Park et al. [2021], neural networks are well known for their ability to learn nonlinear patterns that simpler models may fail to capture and to handle the high dimensionality and sparsity characteristic of microbiome datasets. This capability has been demonstrated on a range of architectures and learning paradigms, from generative adversarial networks to autoencoders, which have shown promise in learning latent representations of microbiome communities that improve phenotype prediction Przymus et al. [2025]. Moreover, DL models are well suited for integrating the multi-modal data typical of microbiome studies. However, most existing DL frameworks rely solely on compositional abundance data, often ignoring valuable microbial metadata such as functional annotations and taxonomic hierarchies, which are critical for capturing biologically meaningful interactions. Incorporating such metadata has been shown to significantly improve the disease prediction performance of DL models Sharma et al. [2020], Metwally et al. [2019]. For example, state-of-the-art classification performance has been achieved by cladogram-based methods that capture taxonomic relationships either by transforming microbiome profiles into phylogenetically ordered images for convolutional neural networks (CNNs) or by constructing multilayer perceptrons (MLPs) whose connectivity mirrors the structure of the cladogram Shtossel et al. [2023], Jiang et al. [2025]. These advances, however, often come at the cost of labor-intensive preprocessing pipelines. More importantly, current methods are limited in their ability to generate biologically actionable insights due to the coarseness of their interpretability outputs (e.g., Grad-CAM heatmaps), reflecting a broader and well-recognized limitation of DL “black box” models Teng et al. [2022]. As a result, a trade-off remains between predictive performance and interpretability in microbiome modeling, making this an active and important area of research in microbiome data science Pan et al. [2024], Przymus et al. [2025].

Graph neural networks (GNNs) are emerging as a powerful approach for modeling microbiome data at the level of microbial interactions, potentially offering both improved disease prediction and enhanced interpretability. Although early GNN-based microbiome studies have demonstrated performance gains over classical machine learning methods, they often required extensive preprocessing to encode microbiome metadata into the graph structure. For example, some approaches constructed cladogram graphs to capture phylogenetic relationships Shtossel et al. [2023], Sharma et al. [2020], while others built microbial co-occurrence networks from data such as abundance correlations Pan et al. [2024] or microbe-disease associations Jiang et al. [2022]. In contrast, Ruaud et al. [2023] introduced a minimal-preprocessing strategy by representing microbes as nodes with microbe-specific genomic sequence embeddings in a complete graph with uniform edge weights, thereby making minimal assumptions about the underlying graph structure. By representing individual microbes as nodes on a graph, this approach allows GNNs to naturally learn the microbiome’s graph structure (“wiring diagram”) directly from training data in an end-to-end manner using their intrinsic message-passing and aggregation functions. Interestingly, this may enable the model to capture high-order and contextual microbe interactions Scarselli et al. [2008], Przymus et al. [2025] that are readily interpretable in the form of edge attributions using existing GNN interpretation methods Liu et al. [2022]. While this holds promise for significantly more refined interpretation insights on microbiome-disease mechanisms, it has yet to be extended to clinically relevant endpoints, where metadata is often heterogeneous and incomplete Kumar et al. [2024]. For example, taxonomic metadata frequently contains gaps at lower taxonomic levels Zhang et al. [2023], but is well known to enhance microbiome-disease prediction Papoutsoglou et al. [2023].

To address these limitations, we introduce GIM (Graph neural network for Interpretable Microbiome), a novel graph-based microbiome modeling and interpretation framework that is end-to-end data-driven (avoids reliance on hand-crafted features) and highly interpretable, without sacrificing predictive performance. GIM models microbial taxonomy as unstructured, sparse node embeddings integrated into an unweighted complete graph. This design allows the GNN to consider every possible pairwise interaction in the community, effectively capturing high-order relationships among nodes, such as multi-node dependencies and network motifs, which extend beyond simple co-occurrence patterns. Importantly, each microbe’s position in the taxonomic hierarchy is encoded into a corresponding node’s feature vector, which injects prior biological knowledge (phylogenetic relatedness) directly into the model’s learning process. Using message passing, GIM learns microbial interaction patterns as a function of fine-grained taxonomic sub-features, capturing nuanced relationships driven by specific microbial traits rather than treating each node as an indivisible entity, which also facilitates fine-grained interpretation through post-hoc gradient-based attribution Liu et al. [2022]. We demonstrate the utility of GIM on a case study of pediatric food allergy, an immune-mediated condition strongly associated with early-life gut microbiome perturbations Davis et al. [2024]. Using a cohort of healthy and allergic subjects, we show that GIM achieves state-of-the-art classification performance, while generating granular interpretations to identify specific microbial taxa and microbial drivers that are most relevant to food allergy status, many of which align with known biological findings. Notably, GIM also generates new hypotheses on microbe pair contextual pathogenicity in allergy. GIM represents a step toward closing the interpretability gap in microbiome machine learning Papoutsoglou et al. [2023], enabling robust predictions and biologically meaningful discoveries from complex microbial datasets. We anticipate that GIM will be broadly applicable to other microbiome-related disorders and will facilitate a deeper understanding of how community interactions contribute to health and disease, representing a critical advance toward actionable microbiome therapeutics.

## 2 Methods

### 2.1 Data

#### Dataset

We utilized the merged allergy dataset from Shtossel et al. [2023], which consisted of a cohort of 1,252 gut microbiome samples, comprising 322 milk allergy, 222 peanut allergy, and 263 tree nut allergy cases (including walnut, hazelnut, and mixed nut allergies), alongside 445 non-allergic controls. Each sample included microbial abundance profiles annotated across seven hierarchical taxonomic levels: kingdom (`k_`), phylum (`p_`), class (`c_`), order (`o_`), family (`f_`), genus (`g_`), and species (`s_`). The dataset exhibited high dimensionality and extreme sparsity with at most 5.1% of a total of 8,425 unique taxa present in any given sample.

#### Data Preprocessing

We transformed each sample into a graph structure where individual microbes were represented as nodes with all-to-all bidirectional connectivity (Figure 1 A). Specifically, each pair of nodes was connected by two directional edges going in opposing directions, representing the potential for two-way interactions between any two microbes in a given microbiome sample. The resulting graph size corresponded directly to the number of microbes present in the sample, with all edges assigned a uniform scalar weight of 1. For each node *i*, we developed a 1-dimensional feature vector by concatenating two components: the scalar abundance value of microbe *i* and its taxa profile encoded as a binary vector. The abundance values underwent preprocessing through logarithmic transformation and normalization to a 0 - 1 range, followed by a logit transformation to center the final abundance values around 0. We constructed microbial taxa features by first aggregating all possible taxa names at each taxonomic level, then creating a binary vector where each microbe’s taxa were indicated by 1 at corresponding indices and 0 elsewhere. For instances where a microbe’s taxonomic classification remained unidentified at certain levels (lacking a taxa name following tag prefixes such as `s_`), we incorporated blank tags as potential taxa names within the vector. Each sample was associated with a binary scalar output, where 1 designated an allergic sample and 0 represented a non-allergic sample. We utilized the `torch_geometric` Data object to convert our graphs into Pytorch tensor objects for further processing.

**Figure 1:**
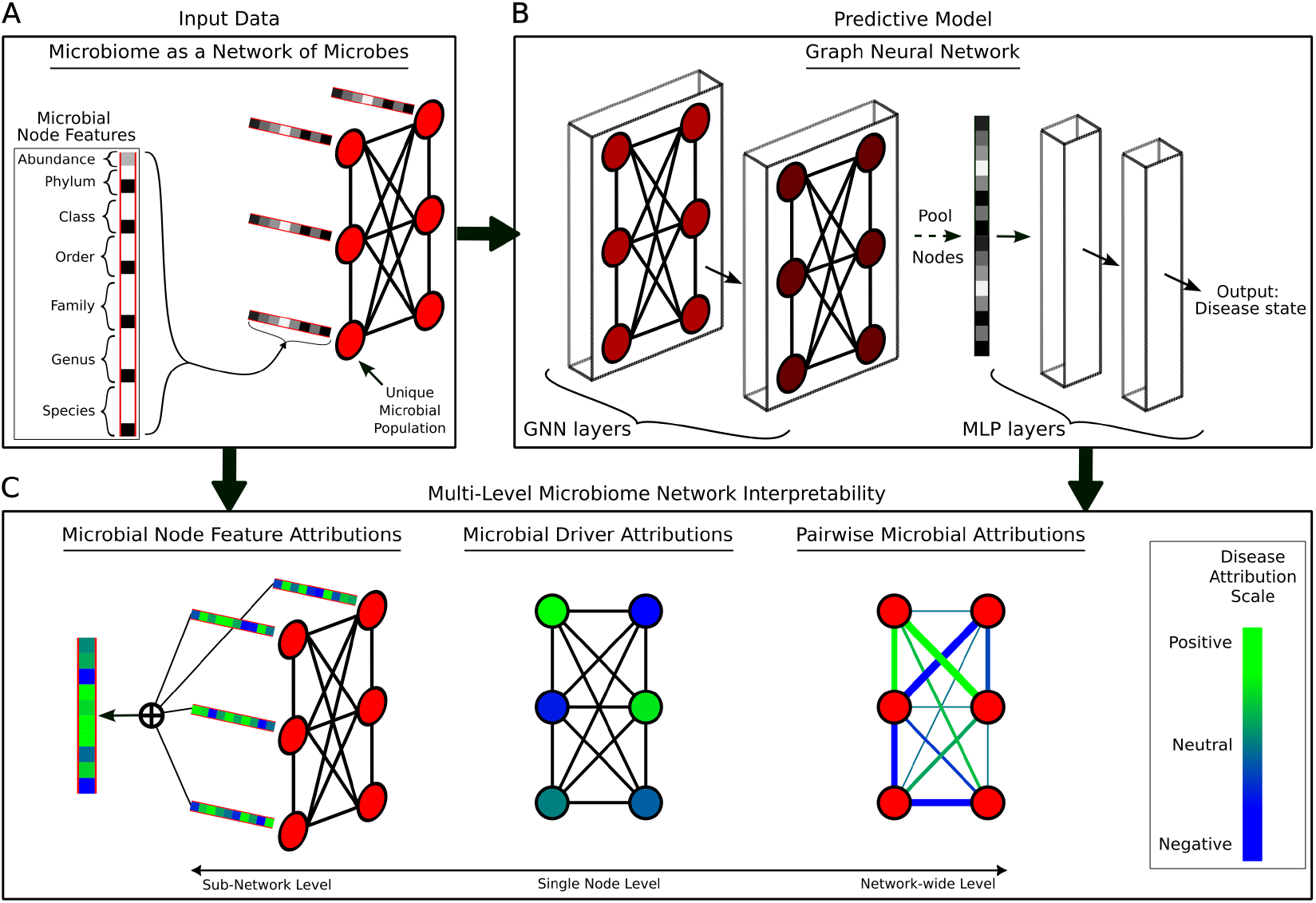
GIM framework for microbiome-disease prediction and interpretability. (**A**) Patient microbiome profiles were modeled as fully connected graphs, where each node corresponds to a microbial population and encodes both its abundance and taxonomic features as a sparse vector representation. These network representations served as input to (**B**) deep predictive models that consisted of graph convolution layers, followed by a node pooling and a sigmoidal output layer. Models predicted the probability of disease, which enabled (**C**) disease specific interpretation of the microbiome network at the (**left**) sub-network level through microbial feature attributions, (**middle**) single node level as microbial driver attributions, and (**right**) network-wide level through pairwise microbial attributions.

### 2.2 Neural Network Training

To model graph-structured data, we developed several GNN architectures, each comprising of *n* graph convolution layers, a node pooling operation, and *m* multi-layer perceptron (MLP) layers (Figure 1 B). Graph convolution layers were implemented using the `SAGEConv` module from Py-Torch Geometric Hamilton et al. [2017]. Node pooling was performed using global mean pooling (`global_mean_pool`) alone or in combination with self-attention graph pooling (`SAGPooling`) Lee et al. [2019] set to selectively retain 80% of nodes based on attention scores. Our baseline model, GIM_1*MP*1_, consisted of a single `SAGEConv` layer transforming input node features from 626 dimensions to 200, followed by global mean pooling to aggregate node representations into a single graph-level vector. This vector was passed through a single-layer MLP with a sigmoid activation function to predict the probability of a given allergy. Further, we tested two variations of the baseline: GIM_2*MP*1_, which added a second `SAGEConv` layer with a 100-dimensional output, and GIM_1*AMP*1_, which incorporated a `SAGPooling` layer before mean pooling. All models were implemented in PyTorch Geometric and parameters were tuned based on the model’s predictive performance and consistency of its feature attributions (See 2.3).

#### Network Training

For optimization, we used the Adam optimizer with a learning rate of 5 × 10^−4^, weight decay regularization of *λ* = 1 × 10^−6^, and a batch size of 2 Kingma and Ba [2017]. To address class imbalance, we implemented weighted binary cross-entropy loss (PyTorch’s `BCEWithLogitsLoss`) where the positive class weight was calculated as the ratio of negative to positive samples. To train models, we used NVIDIA GeForce RTX 2080 Ti GPU.

#### Validation Framework

We implemented a nested k-fold cross-validation strategy with early stopping to facilitate model robustness and prevent overfitting. To ensure comparability with our target performance benchmark Shtossel et al. [2023], we randomly allocated 20% of the dataset to the testing partition of each fold. The validation framework comprised 25 total models trained across 5 outer test folds and 5 nested validation sub-folds. During training of each fold, we continuously monitored validation loss and retained only the best-performing model after seeing no improvement for 25 epochs. In line with Shtossel et al. [2023], we used the weighted area under the curve (AUC) metric to evaluate model performance, with weights accounting for class imbalance in each fold’s test set. All reported AUC scores are the mean and standard deviation across all folds.

### 2.3 Interpretation Methods

We implemented explainable GNN techniques using PyTorch Geometric’s `Explainer` class set to Captum’s Integrated Gradients algorithm Sanchez-Lengeling et al. [2020], Sundararajan et al. [2017]. Our configuration used `node_mask_type=‘attributes’` and `edge_mask_type=‘object’`, with `model_config` parameters set to `mode=‘regression’`, `task_level=‘graph’`, and `return_type=‘raw’`. Despite our model’s binary classification objective, we computed node and edge attributions against raw continuous sigmoidal outputs, which improved interpretation consistency across cross-validation folds.

#### Node Feature Attributions

We calculated node attributions for each sample individually and aggregated the results through three sequential processing steps (Figure 1 C left). First, we normalized the node attributions to unit sum for every sample and applied positive class weighting using the same class weight coefficients employed during GNN training for the respective prediction tasks (milk, peanut, and tree nut allergies), directly aligning the interpretation weights with the original training labels. Next, we summed feature-specific attributions across all samples. Finally, these aggregated attributions were averaged across all folds to ensure robust feature importance quantification (Figure 2 A,C,E).

**Figure 2:**
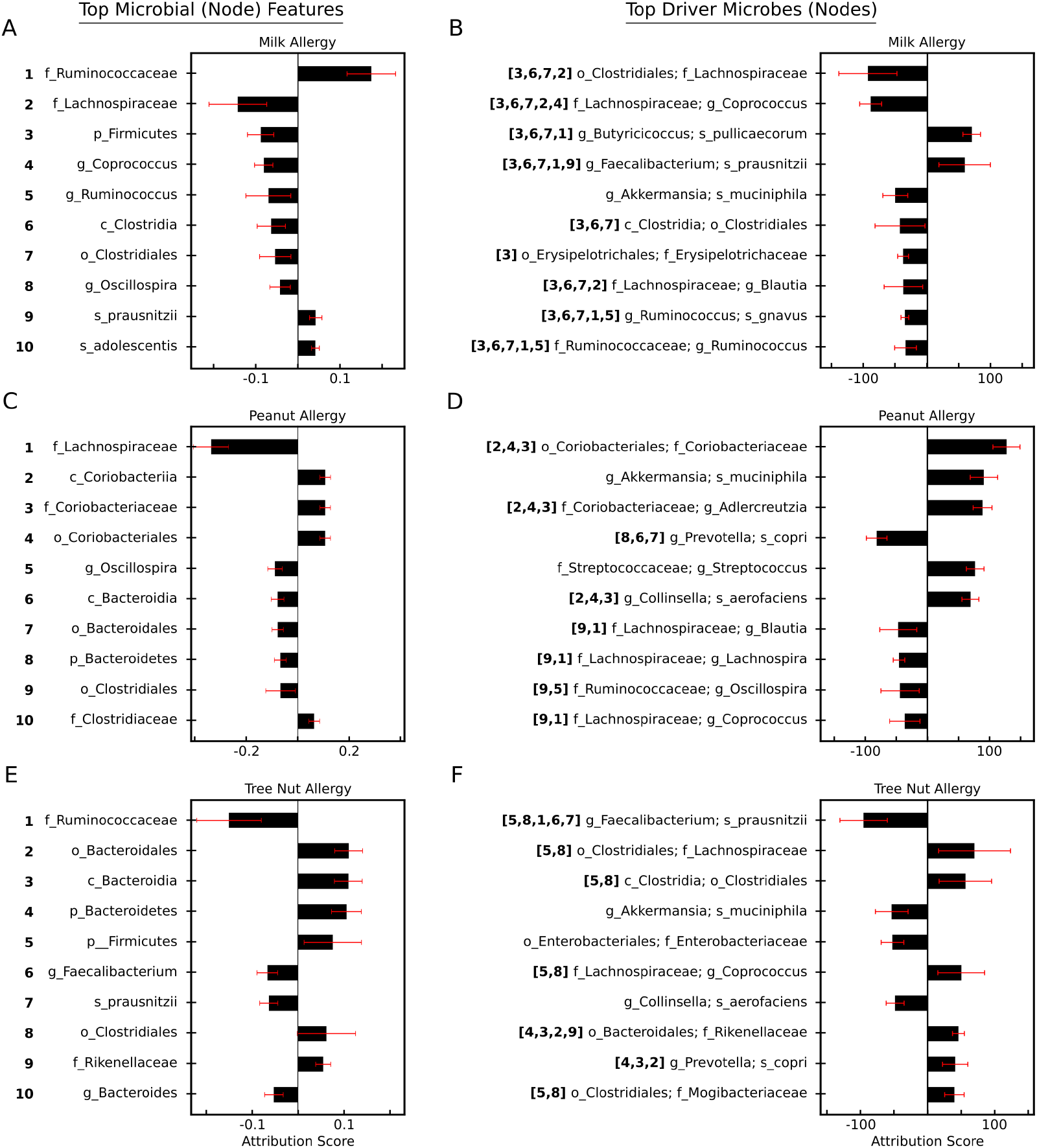
Node feature and microbe driver attributions. Top 10 most influential (**A**,**C**,**E**) microbial taxa based on node features, and (**B**,**D**,**F**) microbial drivers based on edge features. Features/drivers were sorted using attribution magnitude. Taxa features (left) are numbered and mapped to individual microbe drivers (right) using their taxonomic lineage. Bars and error bars represent mean and standard deviation, respectively. (See 2.3)

#### Driver Microbe Attributions

We applied the same normalization procedure and positive class weighting scheme to edge attributions as described above for node feature interpretations (Figure 1 C middle). Following data processing, we aggregated normalized directed edge contributions by source-target pairs across all samples. To quantify the importance of microbial drivers, we leveraged the findings of Tan et al. [2024], which demonstrated that driver microbes can be identified by their outgoing edges in the microbiome causal graph. Accordingly, we computed the sum of attribution scores for all outgoing edges from each source microbe (node) and averaged these scores across all cross-validation folds to derive a measure of driver importance (Figure 2 B,D,F).

#### Dominant Microbe Drivers Analysis

To identify dominant microbial drivers in each sample, we applied a gap-based method to partition driver microbe attributions into high- and low-impact microbial groups. First, microbes were sorted in descending order by their driver attribution scores. The largest gap (i.e., the difference between consecutive scores in the sorted list) was identified, and microbes preceding this gap were classified as dominant drivers. To ensure there was a clear transition between high- and low-impact drivers, gaps were required to be statistical outliers. We defined statistically significant gaps as those exceeding 2 standard deviations from the mean of all observed gaps, with this threshold determined through sensitivity analysis to minimize sample loss while maximizing the number of standard deviations captured (See A.1). Samples that failed this statistical significance test (less than 1% of the dataset) were excluded from downstream analysis. The frequency distribution of dominant drivers across samples (averaged over all cross-validation folds) was visualized after applying a log transformation (with an offset of +2 to stabilize variance) and a rolling average smoothing with a window of 5 (Figure 3 A).

**Figure 3:**
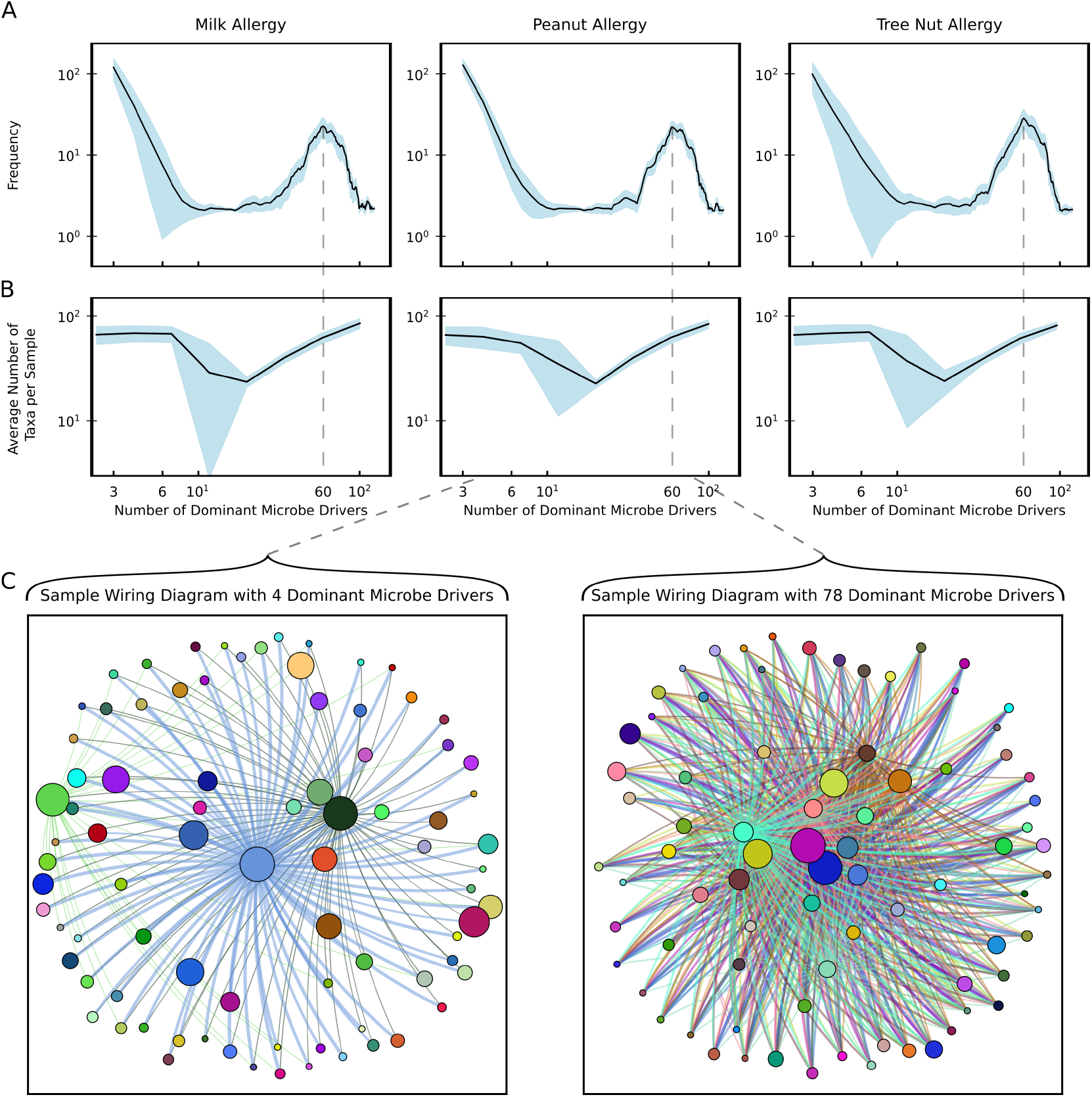
Dominant Microbe Drivers Exhibit a Bimodal Distribution. (A) Frequency distribution and (B) average number of total microbial taxa per sample plotted over the number of dominant driver microbes aggregated across all samples and averaged across all folds. The light blue shaded areas denote standard deviation. (C) Microbiome wiring diagrams for two samples drawn from opposite modes of the bimodal distribution. Visualized edges were filtered to be outgoing from driver microbes and to be larger than threshold magnitudes of 0.2 (left) and 0.4 (right), with magnitudes ranging from 0 to 1. Edge colors match the color of their respective source nodes. Edge width visualizes magnitude of attribution. (See 2.3)

#### Microbe-to-Microbe Attributions

We applied the same normalization procedure and positive class weighting scheme to edge attributions as described above for driver microbes attribution. After processing, we aggregated and summed these normalized edge contributions (ignoring edge direction) for each microbe pair observed in the data’s samples (Figure 1 C right). Finally, we averaged these microbe pair attributions across all folds (Figure 5).

### 2.4 Statistical Methods and Data Presentation

All data in the manuscript are displayed as mean ± standard deviation across cross-validation folds unless specifically indicated. Bar graphs were plotted using Matplotlib library in Python. Network graphs were plotted using Network X library in Python. Specifically, we plotted network graphs in Figure 3 C using `spring_layout` with *k* = 2.2 and in Figure 4 A using `circular_layout`.

**Figure 4:**
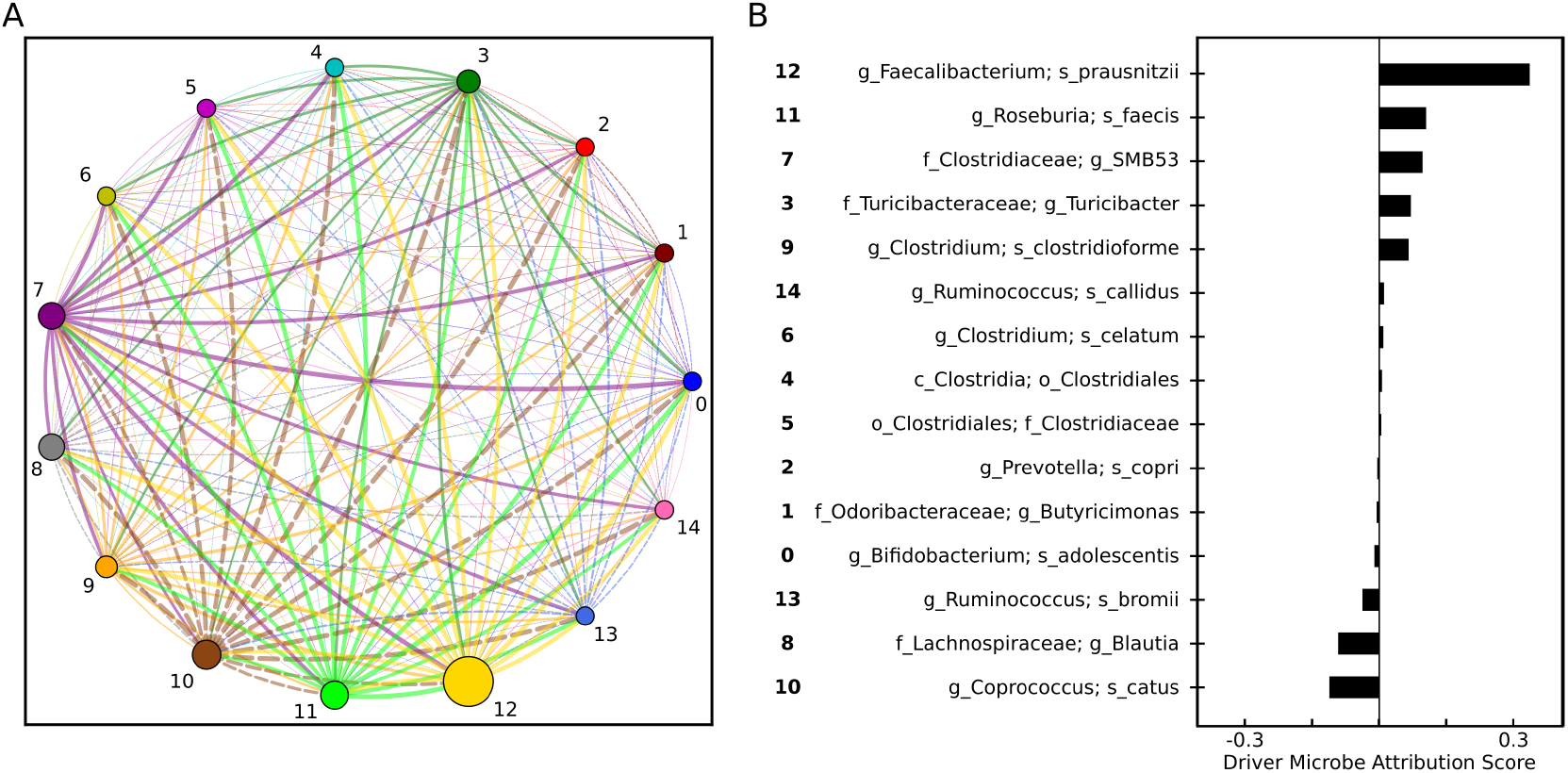
Microbiome wiring diagram and drivers for a single sample. Milk allergy positive sample. (**A**) Sample’s microbiome wiring diagram. Edge color matches the color of its source node. Edge width visualizes magnitude of attribution. Solid/dashed lines denote positive/negative attribution sign. (**B**) Microbe driver attributions for all micrboes in the sample. (See 2.3)

**Figure 5:**
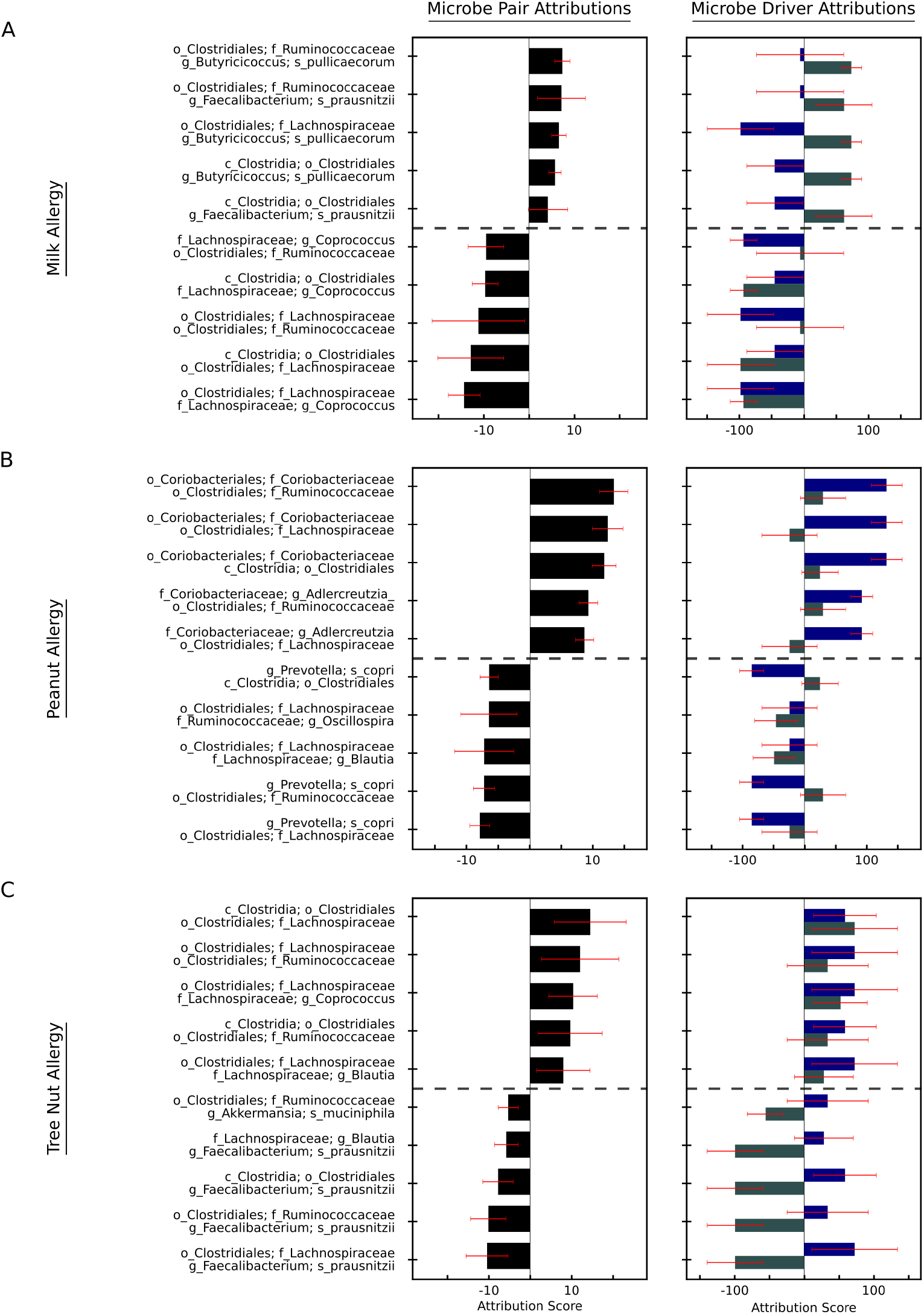
Microbe-to-microbe attributions. (**A**,**B**,**C**) Allergy attributions of top 5 positively and negatively associated microbe pairs (left) along with their corresponding microbe driver breakdown (right). Bars and error bars represent mean and standard deviation, respectively. (See 2.3)

### 2.5 Code and Data Availability

We used data from Shtossel et al. [2023] located at https://github.com/oshritshtossel/iMic/blob/master/Raw_data/Allergy in file `merged_otu_allergy_table_for_Oshrit.txt`. All code used to generate results in this paper can be found at https://github.com/Nexilico/GIM (upon publication).

## 3 Results

### 3.1 End-to-End Representation Learning Outperforms Feature Engineering Methods for Microbiome Disease Prediction

We evaluated GIM’s performance on publicly available microbiome data compiled by Shtossel et al. [2023]. Our baseline GIM model (GIM_1*M*1_) established a new state-of-the-art benchmark with mean AUC values of 0.820 ± 0.024 for milk allergy, 0.777 ± 0.022 for peanut allergy, and 0.705 ± 0.033 for tree nut allergy prediction tasks (Table 1) (Methods 2.2). These results demonstrate substantial improvements over the directly comparable iMic-CNN2-only-micro model, with AUC increases of approximately 0.116, 0.189, and 0.065 on the respective tasks. Notably, our model also outperformed the iMic-CNN2+non-microbial model, which incorporates additional clinical features, by margins of approximately 0.070, 0.144, and 0.057, respectively. This is particularly significant given that GIM relies exclusively on microbial taxonomy and abundance features, highlighting its capacity to capture disease-relevant microbial signals without the need for supplementary clinical data.

**Table 1:**
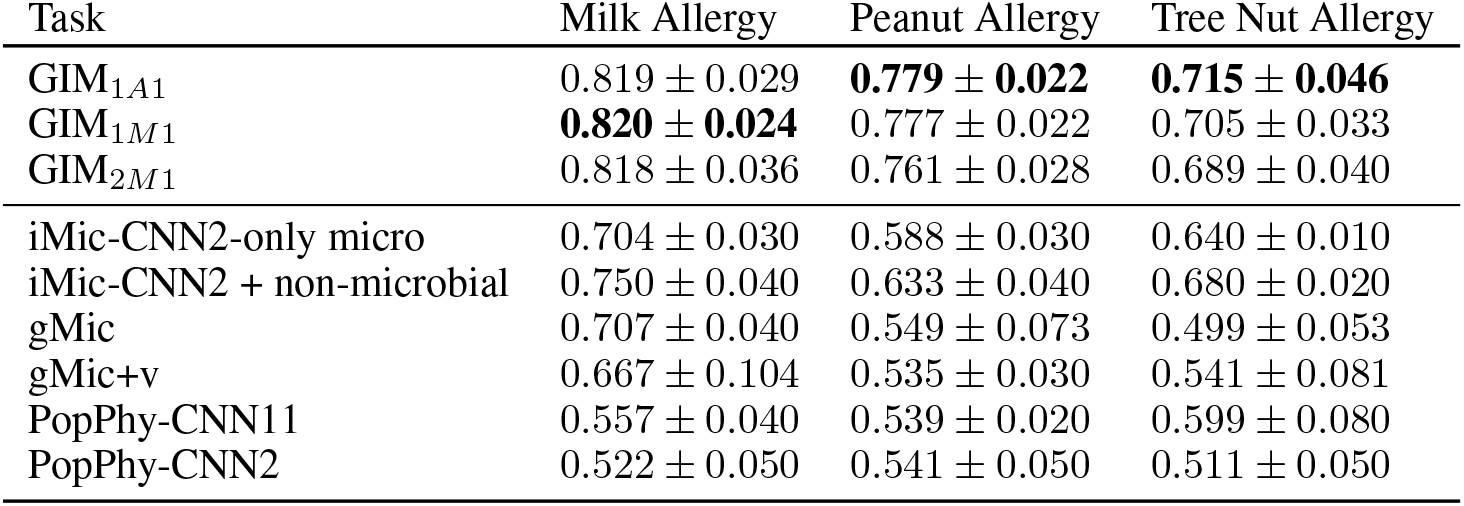
Comparison of model AUC on three binary classification tasks.

Our approach demonstrated robust predictive performance, as evidenced by the minimal variation in AUC scores across our extensive cross-validation testing protocol (Methods 2.2). Specifically, GIM predictions exhibited lower standard deviations compared to most comparable models, with the exception of peanut allergy prediction relative to models iMic-CNN2-only-micro and iMic-CNN2+non-microbial (Table 1). However, the mean AUC performance improvements remained substantial even when this variability was taken into account. Furthermore, our model consistently surpassed previous GNN benchmarks, suggesting that the microbiome feature representation methodology significantly influences model predictive performance.

The improved performance of GIM comes from its graph-based representation of the microbiome that minimizes feature engineering and facilitates end-to-end feature representation learning. While prior GNN benchmarks (e.g. gMic, gMic+v) were trained on microbiome cladogram (taxonomy tree) graphs, our method models microbial interactions through a fully connected graph structure with unweighted bidirectional edges that imposes no a priori assumptions about microbe-to-microbe relationships. Each node in this graph encodes a high-dimensional, sparse feature vector representing a single microbe, with no hierarchical assumptions about its features. The substantial performance gains indicate that this representation more effectively captures the complex dynamics and interactions within microbial communities. This observation suggests that moving away from strictly hierarchical taxonomic representations may better reflect the functional relationships between microorganisms, potentially offering new insights into the underlying biological mechanisms.

We evaluated two variants of our baseline GIM_1*M*1_ model on all three prediction tasks. The GIM_2*M*1_ model, which added a second graph convolution layer to increase depth, showed a modest performance decline, with AUC reductions of −0.002, −0.009, and −0.016 for milk, peanut, and tree nut allergies, respectively, relative to GIM_1*M*1_ (Methods 2.2). This finding suggests that additional graph convolution layers do not enhance performance and may impair it. This is likely due to over-smoothing given our fully connected input graph, which enables each node to aggregate information from all others using a single layer of graph convolution Li et al. [2018]. In contrast, the GIM_1*A*1_ model, which incorporated a global attention-based pooling layer after the graph convolution, yielded varied AUC changes relative to the GIM_1*M*1_ baseline, with gains of 0.002 and 0.010 for peanut and tree nut allergies, respectively, and a reduction of −0.001 for milk allergy. Consequently, we proceeded to perform interpretability analysis using GIM_1*M*1_, given its comparable performance and simpler architecture.

### 3.2 GIM Identifies Biologically Relevant Driver Microbes and Taxa Features

For each prediction task, we computed GIM node feature and edge weight attributions using Integrated Gradients. Node feature attributions reveal which taxa are most positively or negatively associated with a given allergy. However, they do not reveal the taxonomic lineage of specific microbes, because GIM does not use hierarchically structured node features embeddings (Figure 1 A). To identify driver microbes, we analyzed the impact of their outgoing connections on the overall microbiome state Tan et al. [2024] using the computed edge weight attributions (Methods 2.3). We validated our model’s interpretations through two complementary approaches. First, we confirmed that the top attributed taxa and identified driver microbes align with existing literature on known allergy-associated microbes. Second, we examined the overlap between the top taxa features and their presence in top driver microbes, which provides an additional layer of validation for our method (Figure 1 C left, middle).

We identified Ruminococcaceae as the predominant taxon family positively associated with milk allergy, consistent with experimental evidence demonstrating elevated abundances of this microbial family in cow’s milk allergy (CMA) patients Moriki et al. [2024]. Conversely, phylum Firmicutes emerged as a key taxon displaying a negative association with milk allergy, aligning with prior studies reporting reduced Firmicutes abundance in allergic individuals compared to non-allergic, milk-sensitized counterparts Mennini et al. [2021]. This inverse relationship is further supported by clinical evidence linking Firmicutes enrichment to the resolution of milk allergy in pediatric cohorts (Figure 2 A) Bunyavanich et al. [2016], Dong et al. [2018], Yu et al. [2023]. Additionally, our model identified `f_`L. `g_`Blautia and `f_`L. `g_`Coprococcus as putative negative drivers of milk allergy within the gut microbiome, consistent with experimental data implicating depletion of these butyrate-producing genera in allergic states (Figure 2 B) Berni Canani et al. [2016]. In contrast, `f_`F. `g_`Prausnitzii was identified as a positive driver, inline with experimental evidence showing its population enrichment in CMA Xu et al. [2025].

Model interpretation identified Lachnospiraceae as a predominant taxon family negatively associated with peanut allergy, consistent with experimental findings demonstrating its downregulation in the gut microbiota of peanut-allergic murine models Gu et al. [2022]. Notably, Bacteroidales (order), Bacteroidia (class), and Bacteroidetes (phylum) were collectively highlighted as key negatively associated taxa in our model interpretations(Figure 2 C), a pattern further exemplified by the identification of P. Copri as a top negative microbial driver for peanut allergy (Figure 2 D). This observation aligns with clinical evidence showing diminished P. Copri abundance in peanut-allergic patients Goldberg et al. [2020] and, more broadly, reductions in Bacteroidetes populations in allergic individuals Li et al. [2025]. Conversely, Coriobacteriaceae (family), Coriobacteriales (order), and Coriobacteriia (class) all demonstrated positive attribution to peanut allergy and mirroring multiple identified positive driver microbes, which was in line with experimental evidence indicating elevated abundance of Actinobacteria (phylum that encompasses class Coriobacteriia) in allergic individuals Li et al. [2025]. Additionally, our method corroborated L. Blautia as a putative negative driver Li et al. [2025] and A. Muciniphila as a positive driver of peanut allergy, consistent with recent experiemntal studies Parrish et al. [2023].

Interestingly, in contrast to peanut allergy, our model identified P. Copri as a putative positive microbial driver of tree nut allergy, which is supported by experimental evidence demonstrating increased P. Copri abundance in tree nut allergic patients Goldberg et al. [2020]. Forming the taxonomic lineage of P. Copri, Bacteroidales (order), Bacteroidia (class), and Bacteroidetes (phylum) also emerged as top node features with a positive association, some of which were observed experimentally to have elevated populations in patients with tree nut allergy (Figure 2 E,F) Hua et al. [2016].

We observed a bimodal distribution in the number of dominant driver microbes present within individual microbiome samples. Specifically, most samples contained either fewer than 10 dominant drivers or a substantially larger set, typically around 60 (Figure 3 A). Notably, samples with fewer than 10 dominant drivers did not necessarily reflect reduced microbial diversity (Figure 3 B,C). For instance, samples with only three dominant drivers still contained a mean of 70 distinct microbial taxa, suggesting that community influence is concentrated in a small subset of microbes rather than being proportional to overall diversity.

While our model’s aggregated interpretations provide broad insights across the dataset, per-sample analysis remains critical to identify sample-specific microbial drivers obscured by population-level averaging. Examination of a representative milk-allergic sample confirmed the persistence of the above established drivers, including F. Prausnitzii and L. Blautia (Figure 4 B). However, this granular approach also revealed novel candidate drivers such as C. Catus, whose negative association is in line with experimental evidence in milk-allergic cohorts Moriki et al. [2024].

We observed substantial overlap between taxa features and taxonomic lineages of driver microbes, indicating consistency between the node and edge attribution methods, respectively. Specifically, 80%, 100%, and 90% of the top 10 taxa for milk, peanut, and tree nut allergies, respectively, were present in the taxonomic lineages of their respective top 10 driver microbes (Figure 2). This alignment, coupled with independent experimental support, validates that both interpretation methods identify biologically relevant microbial features, reinforcing the robustness of our interpretability framework.

### 3.3 GIM Suggests Existence of Contextual Pathogenicity in the Gut Microbiome

To further demonstrate the interpretive potential of GIM, we show that our model not only identifies individual taxa and driver microbes, but also highlights potential interactions between microbe pairs that may influence disease outcomes (Figure 1 C right). Given that microbial interactions are often highly nonlinear and context-dependent, accurately capturing these relationships remains a significant challenge. To address this, we analyzed pairwise microbial connections using the computed edge weight attributions to assess their impact on the overall microbiome disease state(See 2.3).

Our model identified several instances of putative contextual pathogenicity in both milk and tree nut allergies that would not have been apparent from single-microbe analyses alone. Here, we define contextual pathogenicity as a scenario in which the interaction between two microbes demonstrates a positive or negative association with a given allergy, even when one microbe acts as a positive driver and the other as a negative driver for the same allergy. In such cases, the presence of one microbe appears to modulate, or even reverse, the effect of the other on disease risk. For example, while C. Lachnospiraceae was identified as the top negative driver and B. Pullicaecorum as the top positive driver of milk allergy, their interaction was among the most positively associated with milk allergy (Figure 5 A). Similarly, the interaction between C. Lachnospiraceae and F. Prausnitzii was identified as one of the most negatively associated with respect to tree nut allergy, despite C. Lachnospiraceae being a leading positive driver for that allergy (Figure 5 C). These findings suggest that the impact of C. Lachnospiraceae on allergy risk may depend on the presence or absence of other microbes within the gut microbiome. Such context-dependent effects have been broadly reported in gut microbiome research Khan et al. [2021].

While these findings do not constitute definitive evidence of microbe-to-mictobe effects on allergy, they highlight the model’s capacity to generate testable hypotheses regarding contextual pathogenicity. Future experimental studies will be required to validate these predicted interactions and further elucidate their underlying biological mechanisms, potentially advancing our understanding of the complex microbial dynamics that influence diseases.

## 4 Discussion

In this work, we introduced a microbiome modeling framework, i.e. GIM, that advances the state-of-the-art in both disease prediction performance and fine-grained biological interpretability. Our model generates disease-related mechanistic hypotheses at multiple levels, including individual microbial features, key driver microbes, and specific microbial interactions. Our findings demonstrate that these model-derived insights are consistent with known biological findings, underscoring the potential of GIM to guide and prioritize future experimental studies in microbiome research.

We demonstrated GIM’s robustness to both feature heterogeneity and incompleteness, challenges that are endemic to microbial datasets Kumar et al. [2024]. While we validated GIM’s ability to handle microbial data heterogeneity using both continuous abundance and binary taxa features, the absence of strong structural assumptions in our node encoding makes the method readily extensible to a range of microbial data modalities, including metatranscriptomic, metabolomic, and metaproteomic features. Moreover, GIM’s tolerance for incomplete features, such as missing taxa in our experiments, addresses a key challenge in multi-omics integration, where gaps in data (e.g., incomplete taxa annotations in 16S studies) often limit model generalization and performance Zhang et al. [2023]. Future work could leverage these architectural advantages to unify taxonomic, functional, and clinical metadata, advancing the development of systems-level models for host–microbiome interactions.

By making minimal assumptions about node feature structure and inter-node relationships, GIM’s graph-based microbiome modeling approach naturally extends beyond bacterial taxa to incorporate non-bacterial microbiota, including bacteriophages, fungi, and archaea. This flexibility is essential for accurately modeling the complex, cross-domain interactions within the gut ecosystem, where bacteria, yeasts, viruses, and host factors (such as immune cells, epithelial barriers, and metabolites) collectively shape microbiota-disease relationships. For example, bacteriophages modulate bacterial populations through predation Zuppi et al. [2022], while fungal species like Candida albicans influence mucosal immunity Carlson et al. [2023], underscoring the need for multi-kingdom network analyses. Incorporating these diverse interactions could advance our mechanistic understanding of the pathways linking microbiome dysbiosis to human disease.

The granular interpretability of our approach positions it as a valuable tool for generating testable hypotheses that can guide future microbiome research. By generating attributions for microbial features and identifying microbial drivers, our approach provides detailed insights into influential factors shaping microbiome-associated outcomes. Additionally, microbe-to-microbe attributions can reveal non-trivial interactions, such as contextual pathogenicity. While we verified the biological plausibility of some of the highly attributed mechanisms by cross-referencing with existing literature, these attributions only suggest potential mechanisms of action underlying model predictions and do not establish causality. Experimental validation of contextual pathogenicity hypotheses remains essential, as causality can only be confirmed through rigorous biological studies, which are beyond the scope of this paper. Therefore, any mechanisms suggested by our model should be viewed as a hypothesis to guide and prioritize future experimental work, rather than definitive conclusions.

The GIM framework represents a substantial advancement in microbiome research by enabling the seamless integration of multi-modal microbial data into node vectors within an extensible graph-based representation that can also accommodate non-bacterial microbiome components such as fungi and viruses. Unlike current correlation-based networks or cladogram-based CNN and GNN approaches Shtossel et al. [2023], Jiang et al. [2025], which face challenges in integrating preprocessing-intensive representations with other data modalities and yield only diagnostic associations, GIM explicitly models both bacteria and other components as interacting nodes, thereby capturing relational dynamics through directed edges and offering interpretable insights into microbiome drivers and contextual associations relevant to specific clinical outcomes. This explicit modeling confers unprecedented transparency into the microbiome’s “black box” and, in turn, facilitates therapeutic hypothesis generation, as we demonstrated with GIM_1*M*1_’s extensive interpretability results and robust, state-of-the-art predictive performance. Collectively, these strengths pave the way for future research to expand the GIM framework by incorporating additional multi-modal functional data and non-bacterial entities, and by validating its capacity to elucidate disease-relevant mechanisms and potentially causal interactions, thereby representing a transformative step forward for microbiome therapeutics.

## Acknowledgements

We gratefully acknowledge Dr. Mohammad Mofrad and Dr. Shoko Iwai for their thoughtful feedback and valuable suggestions, which greatly enhanced the clarity and quality of this manuscript. This work is supported by the National Science Foundation (NSF) under Grant No. 2228069.

## A Appendices

### A.1 Threshold Selection for Dominant Driver Microbe Analysis

To determine a statistical threshold for partitioning driver microbes into high- and low-impact groups based on a scaling factor applied to each sample’s gap standard deviation (See 2.3), we conducted a sensitivity analysis. This analysis evaluated the percentage of data samples with largest gaps exceeding the threshold (i.e., statistically significant) as a function of standard deviation scaling factor. We observed that over 99% of samples exhibited their largest gap at more than 2 standard deviations from the mean gap, as shown in Figure 6. Adopting this threshold ensured that over 99% of samples were retained in dominant driver analysis (Figure 3).

**Figure 6:**
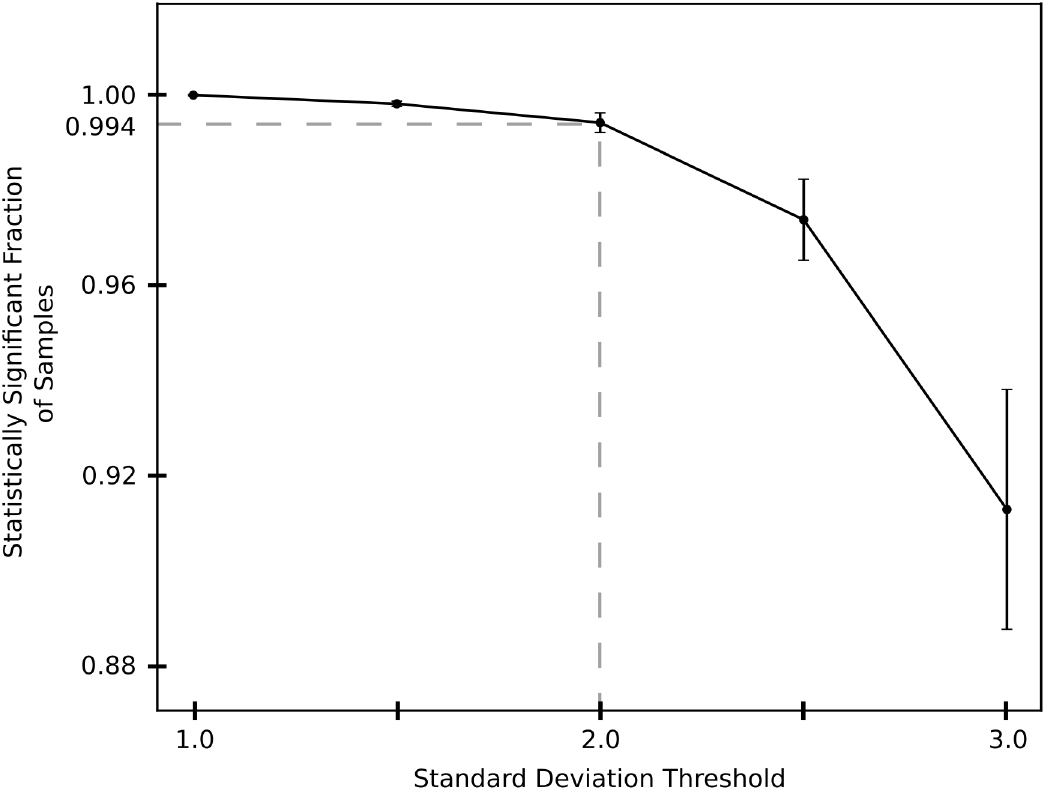
Threshold Sensitivity Analysis. Average fraction of statistically significant samples (greater than threshold) versus standard deviation scaling factor threshold. Fraction of statistically significant samples was computed from driver microbe attributions averaged across all folds and all 3 prediction tasks (milk, peanut, tree nut allergies), with error bars representing standard deviation.

## References

Willem M De Vos, Herbert Tilg, Matthias Van Hul, and Patrice D Cani. Gut microbiome and health: mechanistic insights. Gut, 71(5):1020–1032, 2022.

Yong Fan and Oluf Pedersen. Gut microbiota in human metabolic health and disease. Nature Reviews Microbiology, 19(1):55–71, 2021.

Xuan Zhang, Bei-di Chen, Li-dan Zhao, and Hao Li. The gut microbiota: emerging evidence in autoimmune diseases. Trends in molecular medicine, 26(9):862–873, 2020.

Shokufeh Ghasemian Sorboni, Hanieh Shakeri Moghaddam, Reza Jafarzadeh-Esfehani, and Saman Soleimanpour. A comprehensive review on the role of the gut microbiome in human neurological disorders. Clinical microbiology reviews, 35(1):e00338–20, 2022.

Kerri L Glassner, Bincy P Abraham, and Eamonn MM Quigley. The microbiome and inflammatory bowel disease. Journal of Allergy and Clinical Immunology, 145(1):16–27, 2020.

Thomas CA Hitch, Lindsay J Hall, Sarah Kate Walsh, Gabriel E Leventhal, Emma Slack, Tomas de Wouters, Jens Walter, and Thomas Clavel. Microbiome-based interventions to modulate gut ecology and the immune system. Mucosal immunology, 15(6):1095–1113, 2022.

Huang Lin and Shyamal Das Peddada. Analysis of microbial compositions: a review of normalization and differential abundance analysis. NPJ biofilms and microbiomes, 6(1):60, 2020.

Kevin C Lutz, Shuang Jiang, Michael L Neugent, Nicole J De Nisco, Xiaowei Zhan, and Qiwei Li. A survey of statistical methods for microbiome data analysis. Frontiers in applied mathematics and statistics, 8:884810, 2022.

Jiayu Wu, Kai Wang, Xuemei Wang, Yanli Pang, and Changtao Jiang. The role of the gut microbiome and its metabolites in metabolic diseases. Protein Cell, 12(5):360–373, 12 2020. ISSN 1674-800X. doi: 10.1007/s13238-020-00814-7. URL https://doi.org/10.1007/s13238-020-00814-7.

Stephanie L. Collins, Jonathan G. Stine, Jordan E. Bisanz, C. Denise Okafor, and Andrew D. Patterson. Bile acids and the gut microbiota: metabolic interactions and impacts on disease. Annual Review of Immunology, 21(4):236–247, 2023. ISSN 1740-1534. doi: 10.1038/s41579-022-00805-x.

Ilseung Cho and Martin J Blaser. The human microbiome: at the interface of health and disease. Nature Reviews Genetics, 13(4):260–270, 2012.

Martin Ackermann, Simon van Vliet, et al. Spatial self-organization of metabolism in microbial systems: a matter of enzymes and chemicals. Cell systems, 14(2):98–108, 2023.

Yang-Yu Liu. Controlling the human microbiome. Cell Systems, 14(2), 2023. ISSN 2405-4712. doi: 10.1016/j.cels.2022.12.010.

Xiaoxiu Tan, Feng Xue, Chenhong Zhang, and Tao Wang. mbdriver: identifying driver microbes in microbial communities based on time-series microbiome data. Briefings in Bioinformatics, 25(6): bbae580, 11 2024. ISSN 1477-4054. doi: 10.1093/bib/bbae580. URL https://doi.org/10.1093/bib/bbae580.

Qi Wang, Michael Nute, and Todd Treangen. Bakdrive: Identifying the minimum set of bacterial driver species across multiple microbial communities. bioRxiv, 2021. doi: 10.1101/2021.09.24.461746. URL https://www.biorxiv.org/content/early/2021/09/25/2021.09.24.461746.

Mayank Baranwal, Ryan L Clark, Jaron Thompson, Zeyu Sun, Alfred O Hero, and Ophelia S Venturelli. Recurrent neural networks enable design of multifunctional synthetic human gut microbiome dynamics. eLife, 11:e73870, jun 2022. ISSN 2050-084X. doi: 10.7554/eLife.73870. URL https://doi.org/10.7554/eLife.73870.

Ryan B. Ghannam and Stephen M. Techtmann. Machine learning applications in microbial ecology, human microbiome studies, and environmental monitoring. Computational and Structural Biotechnology Journal, 19, 2021. ISSN 2001-0370. doi: 10.1016/j.csbj.2021.01.028.

Sejun Park, Chulhee Yun, Jaeho Lee, and Jinwoo Shin. Minimum width for universal approximation. In International Conference on Learning Representations, 2021. URL https://openreview.net/forum?id=O-XJwyoIF-k.

Piotr Przymus, Krzysztof Rykaczewski, Adrián Martín-Segura, Jaak Truu, Enrique Carrillo De Santa Pau, Mikhail Kolev, Irina Naskinova, Aleksandra Gruca, Alexia Sampri, Marcus Frohme, et al. Deep learning in microbiome analysis: a comprehensive review of neural network models. Frontiers in Microbiology, 15:1516667, 2025.

Divya Sharma, Andrew D Paterson, and Wei Xu. Taxonn: ensemble of neural networks on stratified microbiome data for disease prediction. Bioinformatics, 36(17):4544–4550, 05 2020. ISSN 1367-4803. doi: 10.1093/bioinformatics/btaa542. URL https://doi.org/10.1093/bioinformatics/btaa542.

Ahmed A Metwally, Philip S Yu, Derek Reiman, Yang Dai, Patricia W Finn, and David L Perkins. Utilizing longitudinal microbiome taxonomic profiles to predict food allergy via long short-term memory networks. PLoS computational biology, 15(2):e1006693, 2019.

Oshrit Shtossel, Haim Isakov, Sondra Turjeman, Omry Koren, and Yoram Louzoun. Ordering taxa in image convolution networks improves microbiome-based machine learning accuracy. Gut Microbes, 15(1):2224474, 2023. doi: 10.1080/19490976.2023.2224474. URL https://doi.org/10.1080/19490976.2023.2224474. PMID: 37345233.

Yifan Jiang, Matthew Aton, Qiyun Zhu, and Yang Young Lu. Modeling microbiome-trait associations with taxonomy-adaptive neural networks. Microbiome, 13, 2025. ISSN 2049-2618. doi: 10.1186/s40168-025-02080-3.

Qiaoying Teng, Zhe Liu, Yuqing Song, Kai Han, and Yang Lu. A survey on the interpretability of deep learning in medical diagnosis. Multimedia Systems, 28(6):2335–2355, 2022.

Shuheng Pan, Xinyi Jiang, and Kai Zhang. Wsgmb: weight signed graph neural network for microbial biomarker identification. Briefings in Bioinformatics, 25(1):bbad448, 2024.

Changzhi Jiang, Minli Tang, Shuting Jin, Wei Huang, and Xiangrong Liu. Kgnmda: A knowledge graph neural network method for predicting microbe-disease associations. IEEE/ACM transactions on computational biology and bioinformatics, 20(2):1147–1155, 2022.

Albane Ruaud, Cansu Sancaktar, Marco Bagatella, Christoph Ratzke, and Georg Martius. Modelling microbial communities with graph neural networks. In NeurIPS 2023 AI for Science Workshop, 2023. URLhttps://openreview.net/forum?id=3Cg94Z1RZj.

Franco Scarselli, Marco Gori, Ah Chung Tsoi, Markus Hagenbuchner, and Gabriele Monfardini. The graph neural network model. IEEE transactions on neural networks, 20(1):61–80, 2008.

Ninghao Liu, Qizhang Feng, and Xia Hu. Interpretability in Graph Neural Networks, pages 121–147. Springer Nature Singapore, Singapore, 2022. ISBN 978-981-16-6054-2. doi: 10.1007/978-981-16-6054-2_7. URL https://doi.org/10.1007/978-981-16-6054-2_7.

Bablu Kumar, Erika Lorusso, Bruno Fosso, and Graziano Pesole. A comprehensive overview of microbiome data in the light of machine learning applications: categorization, accessibility, and future directions. Frontiers in Microbiology, Volume 15 - 2024, 2024. ISSN 1664-302X. doi: 10.3389/fmicb.2024.1343572. URL https://www.frontiersin.org/journals/microbiology/articles/10.3389/fmicb.2024.1343572.

Wenke Zhang, Xiaoqian Fan, Haobo Shi, Jian Li, Mingqian Zhang, Jin Zhao, and Xiaoquan Su. Comprehensive assessment of 16s rrna gene amplicon sequencing for microbiome profiling across multiple habitats. Microbiology Spectrum, 11(3):e00563–23, 2023. doi: 10.1128/spectrum.00563-23. URL https://journals.asm.org/doi/abs/10.1128/spectrum.00563-23.

Georgios Papoutsoglou, Sonia Tarazona, Marta B. Lopes, Thomas Klammsteiner, Eliana Ibrahimi, Julia Eckenberger, Pierfrancesco Novielli, Alberto Tonda, Andrea Simeon, Rajesh Shigdel, Stéphane Béreux, Giacomo Vitali, Sabina Tangaro, Leo Lahti, Andriy Temko, Marcus J. Claesson, and Magali Berland. Machine learning approaches in microbiome research: challenges and best practices. Frontiers in Microbiology, Volume 14 - 2023, 2023. ISSN 1664-302X. doi: 10.3389/fmicb.2023.1261889. URL https://www.frontiersin.org/journals/microbiology/articles/10.3389/fmicb.2023.1261889.

Erin C Davis, Cynthia L Monaco, Richard Insel, and Kirsi M Järvinen. Gut microbiome in the first 1000 days and risk for childhood food allergy. Annals of Allergy, Asthma & Immunology, 133(3): 252–261, 2024.

Will Hamilton, Zhitao Ying, and Jure Leskovec. Inductive representation learning on large graphs. In I. Guyon, U. Von Luxburg, S. Bengio, H. Wallach, R. Fergus, S. Vishwanathan, and R. Garnett, editors, Advances in Neural Information Processing Systems, volume 30. Curran Associates, Inc., 2017. URL https://proceedings.neurips.cc/paper_files/paper/2017/file/5dd9db5e033da9c6fb5ba83c7a7ebea9-Paper.pdf.

Junhyun Lee, Inyeop Lee, and Jaewoo Kang. Self-attention graph pooling. In Proceedings of the 36th International Conference on Machine Learning, 09–15 Jun 2019.

Diederik P. Kingma and Jimmy Ba. Adam: A method for stochastic optimization, 2017. URL https://arxiv.org/abs/1412.6980.

Benjamin Sanchez-Lengeling, Jennifer Wei, Brian Lee, Emily Reif, Peter Wang, Wesley Qian, Kevin McCloskey, Lucy Colwell, and Alexander Wiltschko. Evaluating attribution for graph neural networks. In H. Larochelle, M. Ranzato, R. Hadsell, M.F. Balcan, and H. Lin, editors, Advances in Neural Information Processing Systems, volume 33, pages 5898–5910. Curran Associates, Inc., 2020. URL https://proceedings.neurips.cc/paper_files/paper/2020/file/417fbbf2e9d5a28a855a11894b2e795a-Paper.pdf.

Mukund Sundararajan, Ankur Taly, and Qiqi Yan. Axiomatic attribution for deep networks. In Proceedings of the 34th International Conference on Machine Learning, volume 70, pages 3319–3328. Curran Associates, Inc., 2017. URL https://proceedings.mlr.press/v70/sundararajan17a/sundararajan17a.pdf.

Qimai Li, Zhichao Han, and Xiao-Ming Wu. Deeper insights into graph convolutional networks for semi-supervised learning. In Proceedings of the Thirty-Second AAAI Conference on Artificial Intelligence and Thirtieth Innovative Applications of Artificial Intelligence Conference and Eighth AAAI Symposium on Educational Advances in Artificial Intelligence, AAAI’18/IAAI’18/EAAI’18. AAAI Press, 2018. ISBN 978-1-57735-800-8.

Dafni Moriki, E. Daniel León, Gabriel García-Gamero, Nuria Jiménez-Hernández, Alejandro Artacho, Xavier Pons, Despoina Koumpagioti, Argirios Dinopoulos, Vassiliki Papaevangelou, Kostas N. Priftis, Konstantinos Douros, and M. Pilar Francino. Specific gut microbiome signatures in children with cow’s milk allergy. Nutrients, 16(16), 2024. ISSN 2072-6643. doi: 10.3390/nu16162752. URL https://www.mdpi.com/2072-6643/16/16/2752.

Maurizio Mennini, Sofia Reddel, Federica Del Chierico, Simone Gardini, Andrea Quagliariello, Pamela Vernocchi, Rocco Luigi Valluzzi, Vincenzo Fierro, Carla Riccardi, Tania Napolitano, Alessandro Giovanni Fiocchi, and Lorenza Putignani. Gut microbiota profile in children with ige-mediated cow’s milk allergy and cow’s milk sensitization and probiotic intestinal persistence evaluation. International Journal of Molecular Sciences, 22(4), 2021. ISSN 1422-0067. doi: 10.3390/ijms22041649. URL https://www.mdpi.com/1422-0067/22/4/1649.

Supinda Bunyavanich, Nan Shen, Alexander Grishin, Robert Wood, Wesley Burks, Peter Dawson, Stacie M. Jones, Donald Y.M. Leung, Hugh Sampson, Scott Sicherer, and Jose C. Clemente. Early-life gut microbiome composition and milk allergy resolution. Journal of Allergy and Clinical Immunology, 138(4), 2016. ISSN 0091-6749. doi: 10.1016/j.jaci.2016.03.041.

Ping Dong, Jing jing Feng, Dong yong Yan, Yu jing Lyu, and Xiu Xu. Early-life gut microbiome and cow’s milk allergya prospective case - control 6-month follow-up study. Saudi Journal of Biological Sciences, 25(5):875–880, 2018. ISSN 1319-562X. doi: 10.1016/j.sjbs.2017.11.051. URL https://www.sciencedirect.com/science/article/pii/S1319562X17303248. Genetic characterization of diseases using molecular markers.

Zhidan Yu, Lingling Yue, Zhaojie Yang, Yuesheng Wang, Zihui Wang, Fang Zhou, Chan Li, Lifeng Li, Wancun Zhang, and Xiaoqin Li. Impairment of intestinal barrier associated with the alternation of intestinal flora and its metabolites in cow’s milk protein allergy. Microbial Pathogenesis, 183: 106329, 2023. ISSN 0882-4010. doi: 10.1016/j.micpath.2023.106329. URL https://www.sciencedirect.com/science/article/pii/S0882401023003625.

Roberto Berni Canani, Naseer Sangwan, Andrew T Stefka, Rita Nocerino, Lorella Paparo, Rosita Aitoro, Antonio Calignano, Aly A Khan, Jack A Gilbert, and Cathryn R Nagler. Lactobacillus rhamnosus gg-supplemented formula expands butyrate-producing bacterial strains in food allergic infants. The ISME Journal, 10(3), 2016. ISSN 1751-7370. doi: 10.1038/ismej.2015.151.

Jiaxin Xu, Taha Majid Mahmood Sheikh, Muhammad Shafiq, Muhammad Nadeem Khan, Meimei Wang, Xiaoling Guo, Fen Yao, Qingdong Xie, Zhe Yang, Areeba Khalid, and Xiaoyang Jiao. Exploring the gut microbiota landscape in cow milk protein allergy: Clinical insights and diagnostic implications in pediatric patients. Journal of Dairy Science, 108(1):73–89, 2025. ISSN 0022-0302. doi: 10.3168/jds.2024-25455. URL https://www.sciencedirect.com/science/article/pii/S0022030224011998.

Shimin Gu, Qiang Xie, Chen Chen, Chenglong Liu, and Wentong Xue. Gut microbial signatures associated with peanut allergy in a balb/c mouse model. Foods, 11(10), 2022. ISSN 2304-8158. doi: 10.3390/foods11101395. URL https://www.mdpi.com/2304-8158/11/10/1395.

Michael R. Goldberg, Hadar Mor, Dafna Magid Neriya, Faiga Magzal, Efrat Muller, Michael Y. Appel, Liat Nachshon, Elhanan Borenstein, Snait Tamir, Yoram Louzoun, Ilan Youngster, Arnon Elizur, and Omry Koren. Microbial signature in ige-mediated food allergies. Genome Medicine, 12, 2020. ISSN 1756-994X. doi: 10.1186/s13073-020-00789-4.

Shouming Li, Jingyi Huang, Yunyun Xie, D. Wang, Xin Tan, and Yufan Wang. Investigation of gut microbiota in pediatric patients with peanut allergy in outpatient settings. Frontiers in Pediatrics, Volume 13 - 2025, 2025. ISSN 2296-2360. doi: 10.3389/fped.2025.1509275. URL https://www.frontiersin.org/journals/pediatrics/articles/10.3389/fped.2025.1509275.

Amy Parrish, Marie Boudaud, Erica T. Grant, Stéphanie Willieme, Mareike Neumann, Mathis Wolter, Sophie Z. Craig, Alessandro De Sciscio, Antonio Cosma, Oliver Hunewald, Markus Ollert, and Mahesh S. Desai. Akkermansia muciniphila exacerbates food allergy in fibre-deprived mice. Nature Microbiology, 8, 2023. ISSN 2058-5276. doi: 10.1038/s41564-023-01464-1.

Xing Hua, James J. Goedert, Angela Pu, Guoqin Yu, and Jianxin Shi. Allergy associations with the adult fecal microbiota: Analysis of the american gut project. eBioMedicine, 3, 2016. ISSN 2352-3964. doi: 10.1016/j.ebiom.2015.11.038.

Israr Khan, Yanrui Bai, Lajia Zha, Naeem Ullah, Habib Ullah, Syed Rafiq Hussain Shah, Hui Sun, and Chunjiang Zhang. Mechanism of the gut microbiota colonization resistance and enteric pathogen infection. Frontiers in Cellular and Infection Microbiology, Volume 11 - 2021, 2021. ISSN 2235-2988. doi: 10.3389/fcimb.2021.716299. URL https://www.frontiersin.org/journals/cellular-and-infection-microbiology/articles/10.3389/fcimb.2021.716299.

Michele Zuppi, Heather L. Hendrickson, Justin M. O’Sullivan, and Tommi Vatanen. Phages in the gut ecosystem. Frontiers in Cellular and Infection Microbiology, Volume 11 - 2021, 2022. ISSN 2235-2988. doi: 10.3389/fcimb.2021.822562. URL https://www.frontiersin.org/journals/cellular-and-infection-microbiology/articles/10.3389/fcimb.2021.822562.

Sean L. Carlson, Liya Mathew, Michael Savage, Klaartje Kok, James O. Lindsay, Carol A. Munro, and Neil E. McCarthy. Mucosal immunity to gut fungi in health and inflammatory bowel disease. Journal of Fungi, 9(11), 2023. ISSN 2309-608X. doi: 10.3390/jof9111105. URL https://www.mdpi.com/2309-608X/9/11/1105.

